# Computational pharmacogenomics screen identifies synergistic statin-compound combinations as anti-breast cancer therapies

**DOI:** 10.1101/2020.09.07.286922

**Authors:** Jenna van Leeuwen, Wail Ba-Alawi, Emily Branchard, Joseph Longo, Jennifer Silvester, David W. Cescon, Benjamin Haibe-Kains, Linda Z. Penn, Deena M.A. Gendoo

## Abstract

Statins are a family of FDA-approved cholesterol-lowering drugs that inhibit the rate-limiting enzyme of the metabolic mevalonate pathway, which have been shown to have anti-cancer activity. As therapeutic efficacy is increased when drugs are used in combination, we sought to identify agents, like dipyridamole, that potentiate statin-induced tumor cell death. As an antiplatelet agent dipyridamole will not be suitable for all cancer patients. Thus, we developed an integrative pharmacogenomics pipeline to identify agents that were similar to dipyridamole at the level of drug structure, in vitro sensitivity and molecular perturbation. To enrich for compounds expected to target the mevalonate pathway, we took a pathway-centric approach towards computational selection, which we called mevalonate drug network fusion (MVA-DNF). We validated two of the top ranked compounds, nelfinavir and honokiol and demonstrated that, like dipyridamole, they synergize with fluvastatin to potentiate tumor cell death by blocking the restorative feedback loop. This is achieved by inhibiting activation of the key transcription factor that induces mevalonate pathway gene transcription, sterol regulatory element-binding protein 2 (SREBP2). Mechanistically, the synergistic response of fluvastatin+nelfinavir and fluvastatin+honokiol was associated with similar transcriptomic and proteomic pathways, indicating a similar mechanism of action between nelfinavir and honokiol when combined with fluvastatin. Further analysis identified the canonical epithelial-mesenchymal transition (EMT) gene, E-cadherin as a biomarker of these synergistic responses across a large panel of breast cancer cell lines. Thus, our computational pharmacogenomic approach can identify novel compounds that phenocopy a compound of interest in a pathway-specific manner.

**Significance Statement:** We provide a rapid and cost-effective strategy to expand a class of drugs with a similar phenotype. Our parent compound, dipyridamole, potentiated statin-induced tumor cell death by blocking the statin-triggered restorative feedback response that dampens statins pro-apoptotic activity. To identify compounds with this activity we performed a pharmacogenomic analysis to distinguish agents similar to dipyridamole in terms of structure, cell sensitivity and molecular perturbations. As dipyridamole has many reported activities, we focused our molecular perturbation analysis on the pathway inhibited by statins, the metabolic mevalonate pathway. Our strategy was successful as we validated nelfinavir and honokiol as dipyridamole-like drugs at both the phenotypic and molecular levels. Our pathway-specific pharmacogenomics approach will have broad applicability.

## Background

Triple-negative breast cancer (TNBC) is an aggressive subtype of breast cancer (BC) that has a poorer prognosis amongst the major breast cancer subtypes^1^. This poor prognosis stems from our limited understanding of the underlying biology, the lack of targeted therapeutics, and the associated risk of distant recurrence occurring predominantly in the first two years after diagnosis^2^. Cytotoxic anthracycline and taxane-based chemotherapy regimens remain the primary option for treating TNBC, with other classes of investigational agents in various stages of development. Therefore, novel and effective therapeutics are urgently needed to combat this difficult-to-treat cancer.

Altered cellular metabolism is a hallmark of cancer^3,4^ and targeting key metabolic pathways can provide new anti-cancer therapeutic strategies. Aberrant activation of the metabolic mevalonate (MVA) pathway is a hallmark of many cancers, including TNBC, as the end-products include cholesterol and other non-sterol isoprenoids essential for cellular proliferation and survival^5–7^. The statin family of FDA-approved cholesterol-lowering drugs are potent inhibitors of the rate-limiting enzyme of the MVA pathway, 3-hydroxy-3-methylglutaryl-CoA reductase (HMGCR)^5^. Epidemiological evidence shows that statin-use as a cholesterol control agent is associated with reduced cancer incidence^8^ and recurrence^9–13^. Specifically, in BC, a 30-60% reduction in recurrence is evident amongst statin users, and decreased risk is associated with increased statin duration^9,12,14,15^. We and others have shown preclinically that Estrogen Receptor (ER)-negative BC cell lines, including TNBC, are preferentially sensitive to statin-induced apoptosis^16,17^. Moreover, three preoperative clinical trials investigating lipophilic statins (fluvastatin, atorvastatin) in human BC, showed statin use was associated with reduced tumour cell proliferation and increased apoptosis of high-grade BCs^18,19^. Thus, evidence suggests that statins have potential utility in the treatment of BC, including TNBC.

Drug combinations that overcome resistance mechanisms and maximize efficacy have potential advantages as cancer therapy. Blocking the MVA pathway with statins triggers a restorative feedback response that significantly dampens the pro-apoptotic activity of statins^20,21^. Briefly, statin-induced depletion of intracellular sterols, triggers the inactive cytoplasmic, precursor form of the transcription factor sterol regulatory element-binding protein 2 (SREBP2) to be processed to the active mature nuclear form, which induces transcription of MVA genes, including *HMGCR* and the upstream synthase (*HMGCS1*)^22^. We have shown that inhibiting SREBP2 using RNAi, or blocking SREBP2 processing using the drug dipyridamole, significantly potentiates the ability of statins to trigger tumor cell death^21,23,24^.

Dipyridamole is an FDA-approved antiplatelet agent commonly used for secondary stroke prevention, and since statin-dipyridamole has been co-prescribed for other indications it may be safely used in the treatment of cancer. However, the exact mechanism of dipyridamole action remains unclear as it has been reported to regulate several biological processes. Moreover, the antiplatelet activity of dipyridamole may be a contraindication for some cancer patients. Thus, to expand this dipyridamole-like class of compounds that can potentiate the pro-apoptotic activity of statins, we employed a pathway-centric approach to develop a computational pharmacogenomics pipeline to distinguish compounds that are predicted to behave similarly to dipyridamole in the regulation of MVA pathway genes. Using this strategy, we identified several potential dipyridamole-like compounds including nelfinavir, an FDA-approved antiretroviral drug and honokiol, a compound isolated from *Magnolia spp*., which synergise with statins to drive tumour cell death by blocking the restorative feedback response. Correlation analysis of the statin-compound combination synergy score, with basal mRNA expression across a large panel of BC cell lines, identified *CDH1* expression as a predictive biomarker of response to these combination therapies. Taken together, we provide a new strategy to identify compounds that behave functionally similar to dipyridamole in an MVA pathway-specific manner, and suggest that this approach will have broad utility for compound discovery across a wide-variety of drug/pathway interactions.

## Results

### Computational pharmacogenomic pipeline identifies dipyridamole-like compounds

We developed a computational pipeline that harnesses high-throughput pharmacogenomics analysis to identify dipyridamole-like compounds that synergise with statins by blocking MVA pathway gene expression to inhibit cancer cell viability (**Figure 1**). The LINCS-L1000 (L1000)^25^ and NCI-60^26^ datasets were chosen for these studies as they contain cellular drug-response data at the molecular and proliferative levels across a panel of cell lines, respectively. From these datasets we extracted drug structure, drug-induced gene perturbation data (gene expression changes after drug treatment) and drug-cell line sensitivity profiles for the 238 compounds common to both datasets. Treating each level of data as a separate layer, we restricted the drug-gene perturbation layer from the L1000 dataset to only include the six MVA pathway genes present in the L1000 landmark gene set to enrich for compounds that phenocopy the MVA pathway-specific activity of dipyridamole (**Supplemental Figure 1A**). With dipyridamole as the reference input, we generated an MVA pathway-specific Drug Network Fusion (MVA-DNF) through the integration of 3 distinct data layers: drug structure, MVA-specific drug perturbation signatures, and drug-cell line sensitivity profiles. For each of the data layers incorporated into MVA-DNF, an 238×238 drug affinity matrix was generated, indicating drug similarity for a selected drug against all other drugs. Using the Pearson correlation coefficient, we computed the similarity for every pair of drug perturbation profiles and pairs of drug sensitivity profiles (**Figure 1B**). From this, we identified 23 potential dipyridamole-like compounds that scored as significant (permutation test p-value <0.05); Methods; **Figure 1B and Supplementary Table 1**). Represented as a network, these hits display strong connectivity to dipyridamole as well as to each other.

**Fig. 1.**
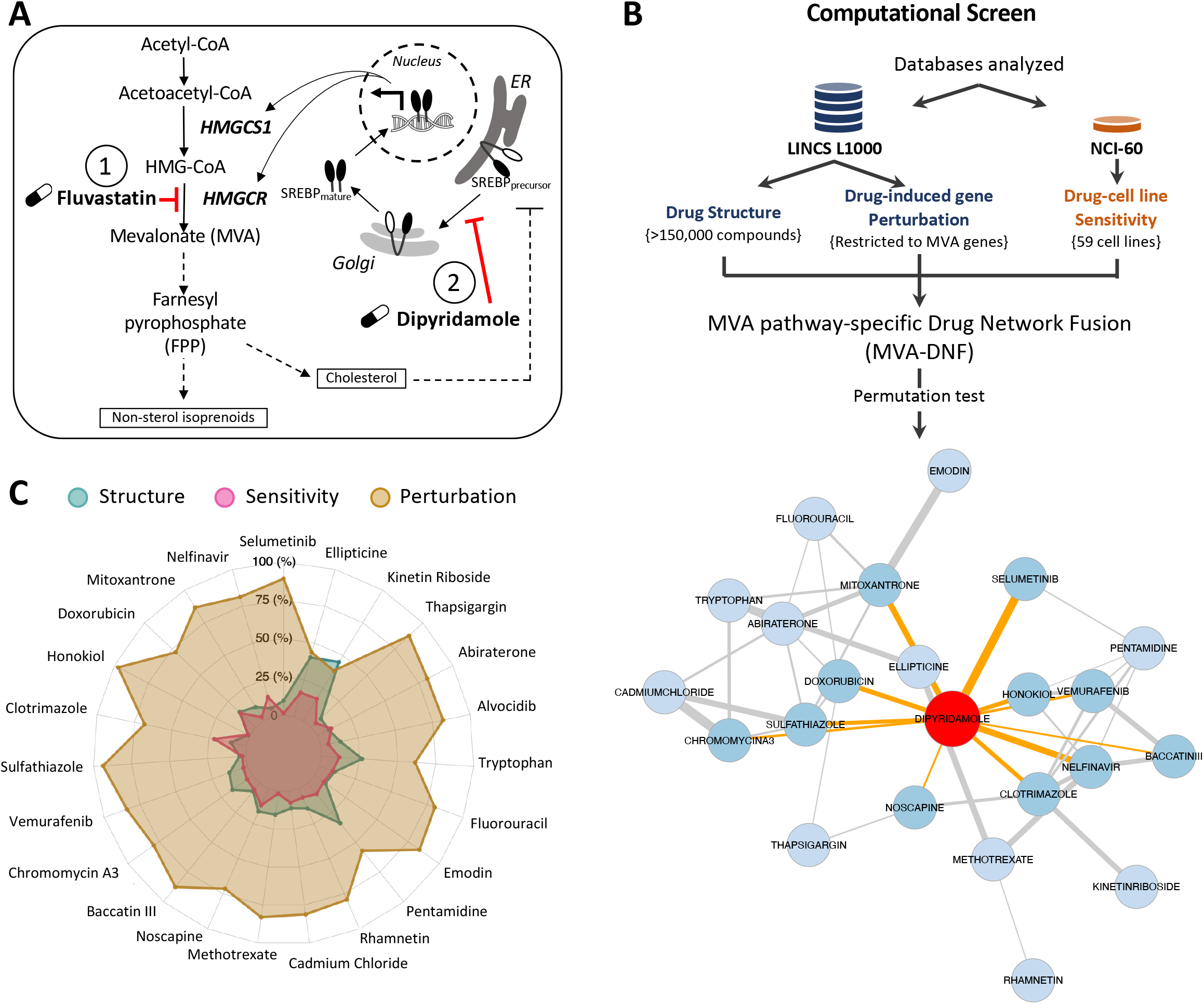
A schematic of the mevalonate (MVA) pathway and overview of the computational pharmacogenomics workflow. **(A)** In response to fluvastatin treatment (labelled with 1), MVA pathway end-product levels decrease, triggering an SREBP-mediated feedback response that activates MVA pathway-associated gene expression to restore cholesterol and other non-sterol end-product levels. Dipyridamole (DP) (labelled with 2) blocks the SREBP-mediated feedback response, thereby potentiating fluvastatin-induced cancer cell death. **(B)** An overview of the computational pharmacogenomics workflow, MVA-DNF, used to identify the top 23 “dipyridamole-like” candidates and visualized as a compound network. MVA-DNF combines drug structure, drug-induced gene perturbation datasets restricted to six MVA pathway-specific genes and drug sensitivity. Permutation specificity testing was performed to select compounds that have a degree of specificity to the mevalonate pathway and dipyridamole. Statistical significance of compounds similar to dipyridamole was assessed by comparing to 1000 networks generated from random selection of six genes within the drug perturbation layer. A network representation of dipyridamole and top 23 statistically-significant (p-value <0.05) “dipyridamole-like” compounds are shown. Each node represents a compound and edges connect compounds based on statistical significance of p-value <0.01. Darker blue nodes and orange edges represent the compounds connected to dipyridamole, and edge thickness represents the associated p-value between the compounds. **(C)** Radar plot of the top 23 dipyridamole-like compounds (p-value <0.05), where the contribution of each individual layer of the MVA-DNF (drug structure, sensitivity, and perturbation) is depicted. Percent contribution of each layer is shown from the center (0%) to the outer edges (100%).

We assessed the contribution of the different data layers (drug structure, drug-gene perturbation, and drug-cell line sensitivity) within the MVA-DNF for each of these 23 compounds (**Figure 1C**). Drug perturbation played a significant role in the selection of novel dipyridamole-like compounds compared to drug sensitivity and drug structure. This reflects the specificity of the MVA-DNF towards the MVA pathway, in comparison to a ‘global’ drug taxonomy that is not MVA pathway-centric. Further assessment of the six MVA-pathway gene expression changes within the drug perturbation signatures highlights comparable expression profiles between dipyridamole and the novel dipyridamole-like compounds (**Supplementary Figure 1B**).

To prioritize and further interrogate the identified dipyridamole-like hits we annotated the 23 compounds by reported mechanism of action and potential clinical utility. Two compounds were excluded from further analysis as they were not clinically useful: Chromomycin A3, a reported toxin^27^, and cadmium chloride, an established carcinogen^28^. The remaining 21 compounds segregated into ten distinct categories, demonstrating that dipyridamole-like hits identified through our pharmacogenomics pipeline spanned a diverse chemical and biological space (**Supplemental Figure 1C, Supplemental Table 1**). We sought to validate the five hits that scored as most similar to dipyridamole, which belong to four different categories (RAF/MEK inhibitor, antiretroviral, anthracycline and natural product). Our lab had previously reported that the anthracycline doxorubicin potentiates lovastatin in ovarian cancer cells^29^ confirming the reliability of our approach. Similarly, RAF/MEK inhibitors such as PD98059 and more recently Selumetinib (AZD6244) have been reported to synergise with statins to potentiate cancer cell death^30,31^. Of the top five hits, doxorubicin was an existing BC chemotherapeutic agent^32^ and therefore removed from further analysis. The molecular targeted compound (selumetinib) along with the novel three compounds were advanced for further evaluation (nelfinavir, mitoxantrone and honokiol) (**Supplemental Table 1**).

### Dipyridamole-like compounds induce apoptosis in combination with fluvastatin and block the sterol-regulated feedback loop of the MVA pathway

To investigate whether the dipyridamole-like compounds could potentiate fluvastatin-induced cell death similar to that of dipyridamole, we first investigated sensitivity to increasing statin exposure in combination with a sub-lethal concentration of the novel dipyridamole-like compounds (**Supplemental Figure 2**) in two breast cancer cell line models with differential sensitivity to fluvastatin as a single agent^16^. As seen with dipyridamole, we observed similar potentiation of fluvastatin (lower IC_50_) when combined with a sub-lethal concentration of selumetinib, nelfinavir, or honokiol, but not mitoxantrone (**Supplemental Fig 3 and Supplemental Fig 4**). Therefore, mitoxantrone was no longer pursued as a dipyridamole-like compound. To determine the nature of the anti-proliferative activity of the statin-compound combinations, we evaluated cell death by fixed propidium iodide staining and PARP cleavage with selumetinib, nelfinavir, or honokiol. Our data indicate that all three compounds, at concentrations that have minimal effects as single agents, phenocopy dipyridamole and potentiate statin-induced cell death (**Figure 2A-C**).

**Fig. 2.**
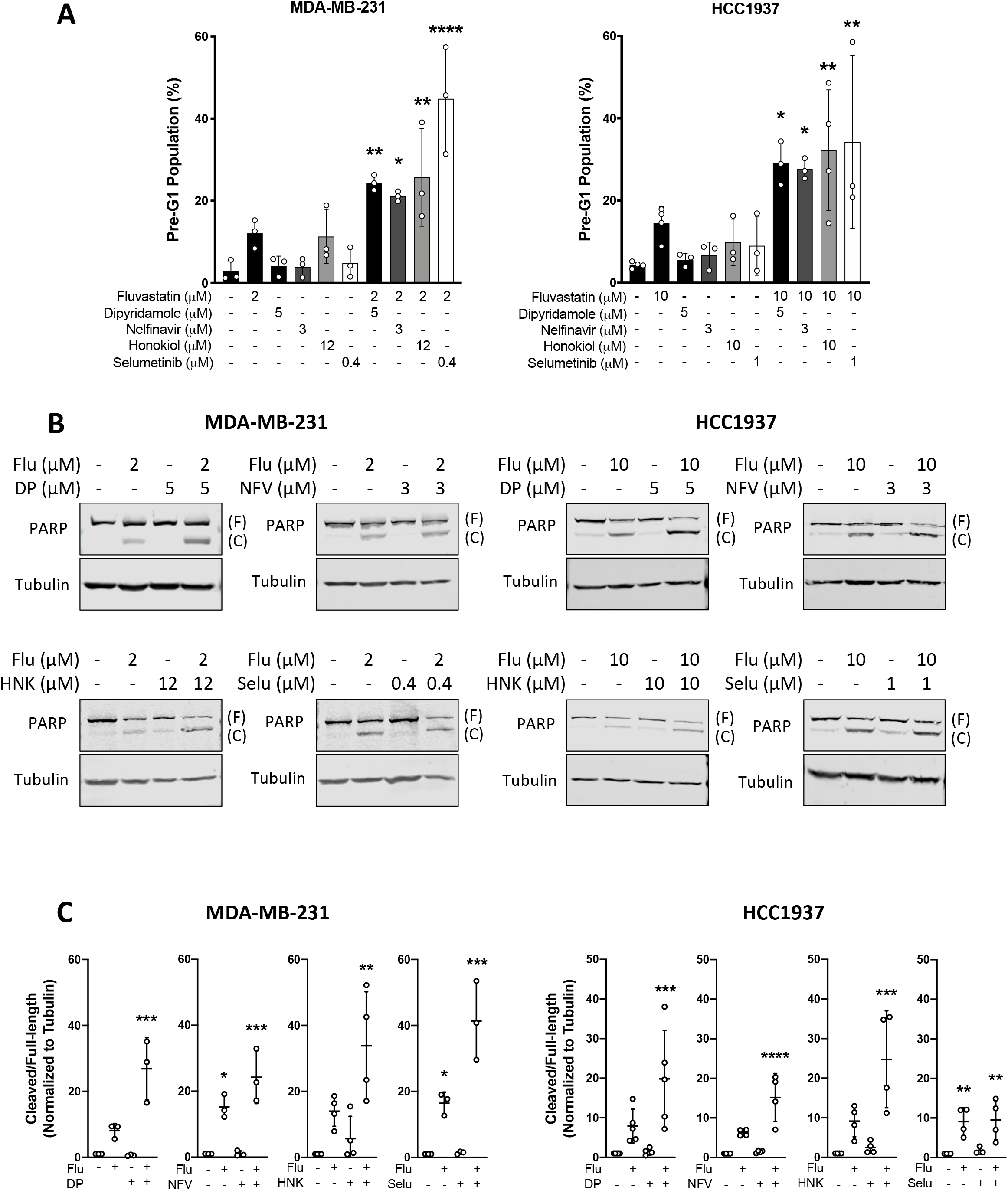
Dipyridamole-like compounds potentiate fluvastatin-induced cell death. **(A)** MDA-MB-231 and HCC1937 cells were treated with solvent controls or fluvastatin +/− dipyridamole (DP), nelfinavir (NFV), honokiol (HNK) or selumetinib (Selu) for 72 hours, fixed in ethanol and assayed for DNA fragmentation (% pre-G1 population) as a marker of cell death by propidium iodide staining. Error bars represent the mean +/− SD, n = 3-4, *p < 0.05, **p < 0.01, ****p < 0.0001 (one-way ANOVA with Bonferroni’s multiple comparisons test, where each treatment was compared to the solvent control). **(B)** Cells were treated as in (A), protein isolated and immunoblotting was performed to assay for PARP cleavage. (F) represents full-length PARP and (C) represents cleaved PARP. **(C)** PARP cleavage (cleaved/full-length) shown in (B) was quantified by densitometry and normalized to Tubulin expression. Error bars represent the mean +/− SD, n = 3-5, *p < 0.05, **p<0.005, ***p<0.001, ****p<0.0001 (one-way ANOVA with Bonferroni’s multiple comparisons test, where each group was compared to the solvent control within each experiment).

Mechanistically, statins induce a feedback response mediated by SREBP2 that has been shown to dampen cancer cell sensitivity to statin exposure. Moreover, blocking the SREBP2-mediated feedback response with dipyridamole enhances statin-induced cancer cell death^21,24^. We have shown that dipyridamole blocks the regulatory cleavage and therefore activation of SREBP2, decreasing mRNA expression of SREBP2-target genes of the MVA pathway. As expected, statin treatment induced the expression of SREBP2-target genes, *INSIG1, HMGCR* and *HMGCS1* after 16 hr of treatment, which was blocked by the co-treatment with dipyridamole (**Figure 3A, Supplemental Figure 5A**). Similarly, nelfinavir and honokiol both phenocopy dipyridamole and block the statin-induced expression of MVA pathway genes (**Figure 3A, Supplemental Figure 5A**). By contrast, co-treatment with selumetinib did not block the fluvastatin-induced feedback response. Housekeeping gene *RPL13A* was used as a reference gene for normalizing mRNA between samples and was not altered in the presence of the compounds (**Supplemental Figure 5B**).

**Fig. 3.**
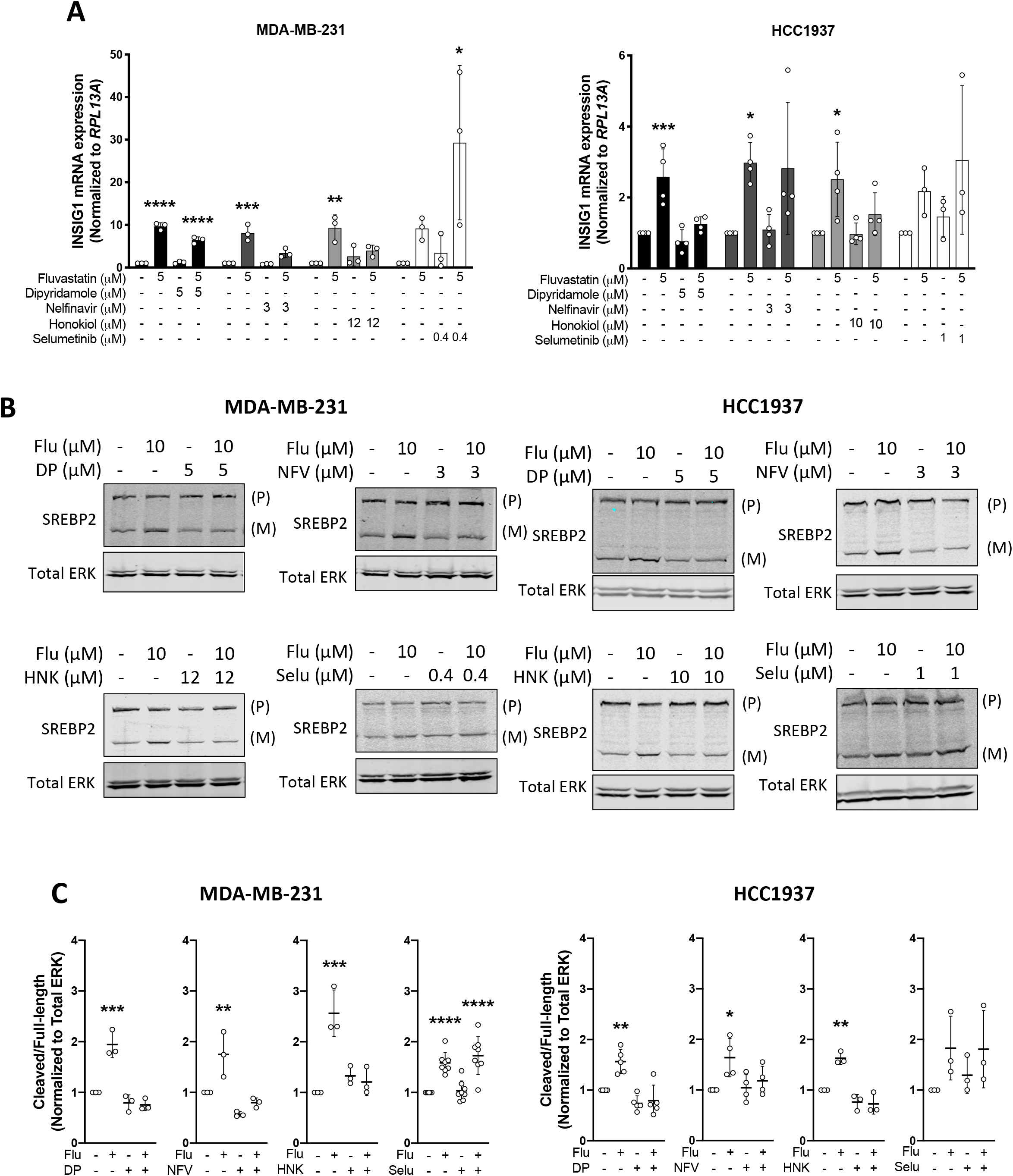
Nelfinavir and Honokiol block fluvastatin-induced SREBP activation. **(A)** MDA-MB-231 and HCC1937 cells were exposed to solvent controls, fluvastatin +/− dipyridamole, nelfinavir, honokiol or selumetinib for 16 hours, and RNA was isolated to assay *INSIG1* expression by qRT-PCR. mRNA expression data are normalized to *RPL13A* expression. Error bars represent the mean +/− SD, n = 3-4, *p < 0.05, **p<0.005, ***p<0.001, ****p<0.0001 (one-way ANOVA with Bonferroni’s multiple comparisons test, where each group was compared to the solvent control group within each experiment). **(B)** MDA-MB-231 and HCC1937 cells were treated with fluvastatin +/− dipyridamole, nelfinavir, honokiol or selumetinib for 12 hours, and protein was harvested to assay for SREBP2 expression and cleavage (activation) by immunoblotting. (P) represents precursor, full-length SREBP2 and (M) represents mature, cleaved SREBP2. **(C)** SREBP2 cleavage (cleaved/full-length) was quantified by densitometry and normalized to total ERK expression. Error bars represent the mean +/− SD, n = 3-8, *p < 0.05, **p<0.005, ***p<0.001, ****p<0.0001 (one-way ANOVA with Bonferroni’s multiple comparisons test, where each group was compared to the solvent controls group within its experiment).

Because SREBP2 is synthesized as an inactive full-length precursor that is activated to the mature nuclear form upon proteolytic cleavage, we used western blot analysis to assess the protein levels of both full-length and mature SREBP2. Nelfinavir and honokiol, but not selumetinib, blocked fluvastatin-induced SREBP2 processing and cleavage similar to that of dipyridamole (**Figure 3B-C**). This suggests that while selumetinib is a strong potentiator of statin induced cell death, it does not mimic the action of dipyridamole by blocking the restorative feedback response (**Figure 3, Supplemental Figure 5**).

### Novel statin-compound combinations phenocopy synergistic activity of fluvastatin-dipyridamole in a breast cancer cell line screen

To investigate whether the potentiation of fluvastatin by nelfinavir and honokiol has broad applicability and examine the determinants of synergy, we further evaluated these statin-compound combinations across a large panel of 47 breast cancer cell lines. A 5-day cytotoxicity assay (sulforhodamine B assay; SRB) in a 6×10 dose matrix was used to assess fluvastatin-compound efficacy. As expected, dipyridamole treatment resulted in a dose-dependent decrease in fluvastatin IC_50_ value (**Supplemental Figure 6A**). Similarly, nelfinavir and honokiol treatment also resulted in a dose-dependent decrease in fluvastatin IC_50_ values similar to that of dipyridamole (**Supplemental Figure 6A**). This suggests that our computational pharmacogenomic pipeline predicts compounds that potentiate statin activity similarly to dipyridamole across multiple subtypes of breast cancer cell lines.

Next we evaluated statin-compound synergy using the Bliss Index model derived using SynergyFinder^33^ across the panel of breast cancer cell lines. Like the dose dependent sensitivity data, we observed that the trend in synergy between fluvastatin-dipyridamole across the 47 breast cancer cell lines was also seen with fluvastatin-nelfinavir and fluvastatin-honokiol (**Figure 4A**). Since we had previously identified that the basal subtype of breast cancer cell lines were more sensitive to single agent fluvastatin^16^, we evaluated whether basal breast cancer cell lines were similarly more sensitive to the fluvastatin-compound combinations. Using the SCMOD2 subtyping scheme, we evaluated the basal, HER2 and luminal B status of each cell line and determined synergy is not dependent on BC subtype (**Supplemental Figure 6B**) suggesting that these statin-compound combinations can be applied to multiple breast cancer subtypes as therapeutic options.

**Fig. 4.**
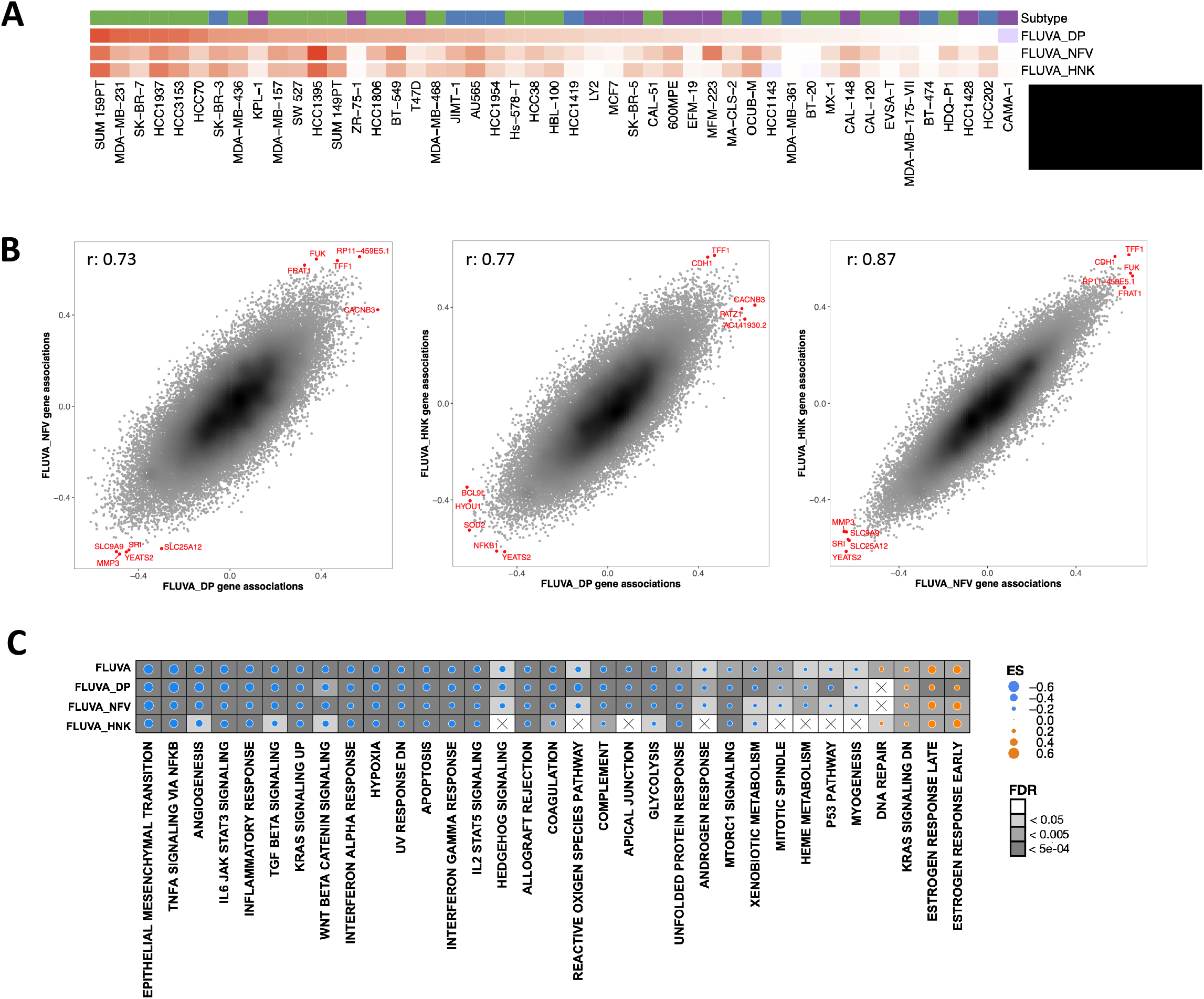
Compound combination synergy analysis. **(A)** Heatmap of synergy scores (Bliss Index model) for fluvastatin (Fluva) + dipyridamole (DP), nelfinavir (NFV) or honokiol (HNK) in a panel of 47 breast cancer cells lines. Ordered by synergy score of Fluva + DP, from greatest to least synergy. Breast cancer subtype of each cell line is shown and is based on the SCMOD2 subtyping scheme. **(B)** Basal mRNA expression^34^ associations with synergy scores between each drug combination (e.g. Fluva+NFV vs. Fluva+DP, Fluva+HNK vs. Fluva+DP, and Fluva+NFV vs. Fluva+HNK). Correlations were calculated using Pearson correlation coefficient. Top five basally-expressed genes associated with synergy in either direction are annotated in red. **(C)** Gene set enrichment analysis (GSEA) using the Hallmark gene set collection, where genes were ranked according to their correlation to the fluvastatin IC_50_ (Fluva) value or to the synergy score (Fluva+DP, Fluva+NFV and F+HNK). Dot plot was restricted to pathways enriched in two out of four groups. Dot size indicates the difference in enrichment scores (ES) of the pathways. Background shading indicates the FDR. X indicates pathway and drug combinations that were not significantly enriched (FDR > 0.05).

Because the synergy profiles across the three fluvastatin-compound combinations were significantly similar, we next interrogated whether baseline gene and/or protein expression profiles across the cell lines for each of the statin-compound combinations was associated with synergy. To further interrogate the similarity between the statin-compound combinations, we correlated the RNA-seq and reverse phase protein array (RPPA) profiles of the 47 breast cancer cell lines^34^ with their synergy scores for each of the statin-compound combinations. These represent the transcriptomic and proteomic state associations with synergy for each combination. We then evaluated the correlation between these associations across the different combinations (F+DP vs F+NFV; F+DP vs F+HNK; F+NFV vs F+HNK) (**Figure 4B**) and found a high positive correlation between the combinations on the basis of similar transcriptomic associations (F+NFV vs F+DP, <u>R=0.73</u>; F+HNK vs F+DP, <u>R=0.77</u>; F+NFV vs F+HNK, <u>R=0.87</u>). This high positive correlation was also seen between these combinations using proteomic (RPPA) and synergy data (**Supplemental Figure 6C**) suggesting that similar pathways were associated with the synergistic response to the three statin-compound combinations.

To compare the overlap in pathways associated with sensitivity to fluvastatin alone, and synergy between the fluvastatin-compound combinations, a Gene Set Enrichment Analysis (GSEA) using the Hallmark Gene Set Collection was performed^35^. These results showed that enriched pathways were highly similar amongst fluvastatin alone and the fluvastatin-compound combinations with one of the highest scoring enriched pathways being EMT (**Figure 4C**). To further support this finding and because of the low agreement amongst EMT gene sets, we also evaluated four additional GSEA EMT pathways and observed similar trends between fluvastatin alone and the fluvastatin-compound combinations for each of the EMT gene sets (**Supplemental Figure 6D**). As we and others have published that mesenchymal-enriched cancer cell lines are more sensitive to statin monotherapy^36,37^, this data suggests that fluvastatin is the primary driver of response to these statin-compound combinations. This is consistent with fluvastatin inhibiting the MVA pathway, triggering the SREBP-mediated feedback response, which in turn is inhibited by the second compound (dipyridamole, nelfinavir or honokiol) in these fluvastatin-compound combinations.

We then examined the individual genes within each of the GSEA EMT pathways to identify a biomarker of synergy to the statin-compound combinations. Within the EMT field, gene set signatures have low agreement (**Supplemental Figure 7**). Previously our lab published a binary classifier of five EMT genes to predict increased sensitivity to statins across 631 cell lines representing multiple cancer types^36^. We evaluated whether this binary five-gene classifier could also predict synergy between the different fluvastatin-compound combinations. The five-gene EMT classifier could predict sensitivity to fluvastatin alone across the panel of breast cancer cell lines (**Supplemental Figure 8A**), but failed to predict synergy to the fluvastatin-compound combinations (**Supplemental Figure 8B**). We next interrogated each of the five genes individually. Interestingly, low gene expression and protein levels of E-cadherin *(CDH1),* a canonical epithelial state marker, not only predicted sensitivity to fluvastatin, but also demonstrated synergy across all three fluvastatin-compound combinations (**Figure 5A-B and Supplemental Figure 8C**). To validate our findings, we probed for basal E-cadherin protein expression across a panel of nine breast cancer cell lines and showed that synergy to the novel statin-compound combinations is positively associated with low E-cadherin protein expression (**Figure 5C-D**). Overall, this data validates that our MVA-DNF pharmacogenomics strategy can successfully distinguish compounds that, like-dipyridamole, can synergize with statins to trigger BC tumour cell death.

**Fig. 5.**
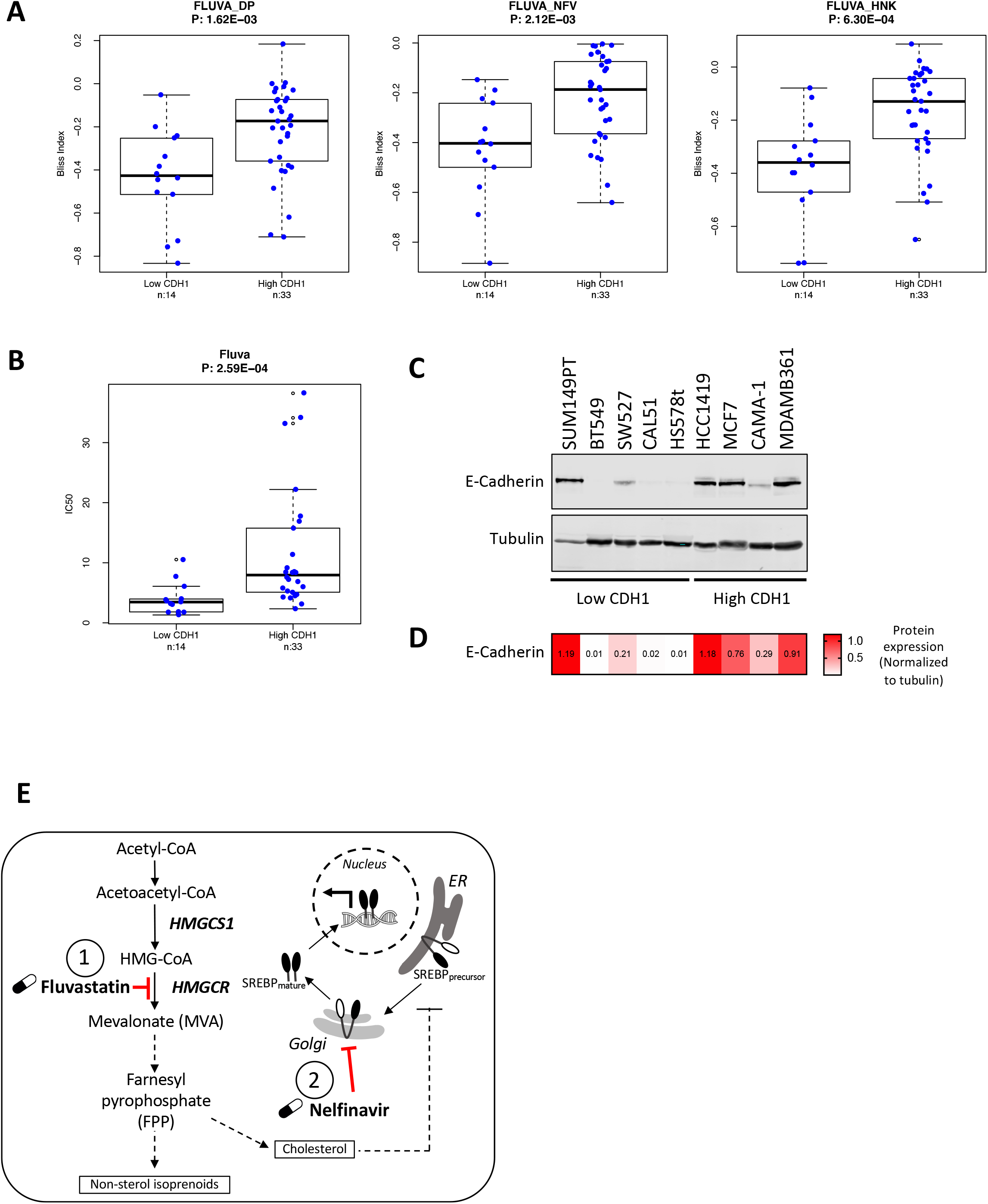
Basal E-cadherin predicts synergistic response to fluvastatin+compound combinations. **(A)** Basal E-cadherin mRNA expression between cell lines predicted to be synergistic or not to each drug combination. Synergy was defined by Bliss Index and significance was measured by wilcoxon rank sum test. **(B)** Basal E-cadherin mRNA expression between cell lines predicted to be respondent or not to fluvastatin. Sensitivity was defined by IC_50_ and significance was measured by wilcoxon rank sum test. **(C)** Protein lysates were isolated from a panel of breast cancer cell lines to assay for basal E-cadherin expression by immunoblotting. **(D)** Densitometry values of normalized E-cadherin expression plotted as a heatmap. E-cadherin expression was quantified by densitometry and normalized individually to Tubulin expression. **(E)** Schematic diagram detailing the potential for fluvastatin (labelled with 1) and nelfinavir (labelled with 2) to block the SREBP2-mediated feedback response and synergize to potentiate fluvastatin-induced cell death.

## Discussion

By blocking the statin-induced restorative feedback response, dipyridamole potentiates statin efficacy to drive tumor cell death^21,24^. However, due to the polypharmacology of dipyridamole and the potential contraindication of this platelet-aggregation inhibitor for some cancer patients, it was essential to identify additional dipyridamole-like compounds and expand this class of agents to provide synergistic statin+compound treatment options for cancer therapy. To this end, we developed a novel computational pharmacogenomics pipeline that distinguished compounds that are similar to dipyridamole at the level of structure, MVA pathway gene expression perturbation, and anti-proliferative activity. We identified 23 potential dipyridamole-like compounds and then evaluated several of the top hits for their ability to phenocopy dipyridamole. Through this approach, we validated that nelfinavir and honokiol sensitize breast cancer cell lines to statin-induced cell death by blocking the statin-induced restorative feedback loop. Analysis of basal RNA and protein expression identified the canonical EMT gene *CDH1* (E-cadherin) as a biomarker of the synergistic response to both statin+nelfinavir and statin+honokiol treatment. Thus, the computational pharmacogenomics screen described here identified synergistic statin-compound drug combinations as novel anti-breast cancer therapies.

The integration of a computational pharmacogenomics pipeline and cellular validation to identify novel compounds with similar biological activities provides a rapid and inexpensive strategy that has potential broad applicability as it is also adaptable. For example, one issue we had to overcome in identifying dipyridamole-like compounds was the polypharmacology of dipyridamole itself. Dipyridamole was originally identified for its anti-platelet aggregation activity and thus the mechanism of action remains unclear. Several activities of dipyridamole have been described including an inhibitor of phosphodiesterases (PDEs)^38^, nucleoside transport^39^ and glucose uptake^40^. The complexity associated with this polypharmacological activity beyond the mevalonate pathway was circumvented by restricting the gene perturbation layer of the DNF to MVA pathway genes. This shows that the computational pharmacogenomics pipeline described here is likely tunable to drug-specific structural features, activities and signaling pathways.

The new statin-sensitizing agents identified here using MVA-DNF include nelfinavir and honokiol, which like dipyridamole, inhibit statin-induced SREBP2 cleavage and activation^21,24^. To date, a number of SREBP2 inhibitors have been identified that block SREBP2 processing from its precursor to mature form, including fatostatin, betulin, and xanthohumal (ER-Golgi translocation), PF-429242 (site-1 protease (S1P) cleavage), and nelfinavir and 1,10-phenanthroline (site-2 protease (S2P) cleavage). Additional SREBP2 inhibitors include BF175 and tocotrienols that target SREBP2 transcriptional activity and protein stability, respectively. However, other than nelfinavir, these agents have many reported targets and are only used as tool compounds for research purposes.

The S2P protease inhibitor nelfinavir was approved for use in 1997 as an antiviral for the treatment of HIV, and in recent years has begun to be evaluated for its utility as an anti-cancer agent^41^. While combination studies of statins and nelfinavir have not been previously reported or investigated in the context of cancer, open-label, multiple-dose studies have been performed to determine the interactions between nelfinavir and two statins (atorvastatin and simvastatin) in healthy volunteers. It was stated that co-administration of nelfinavir and simvastatin should be avoided while atorvastatin should be co-administered with caution. It should be noted that the family of statin drugs are metabolized by different enzymes. Therefore, these interactions of nelfinavir with atorvastatin and simvastatin were likely due to drug-drug interactions leading to the inhibition of CYP3A4. By contrast, fluvastatin is metabolized by CYP2C9 providing additional rationale for our use of fluvastatin in statin-drug combinations as the probability of drug-drug interactions is significantly reduced.

To the best of our knowledge, this is the first study to report honokiol to synergize with statins in the context of cancer. Honokiol is a natural product commonly used in traditional medicine and has a number of reported mechanisms of action. How honokiol inhibits SREBP2 remains unknown, however this is the first study to interrogate its activity in SREBP2 translocation and gene expression alone and in combination with statins. As honokiol and its derivatives are presently under development, these data can now be incorporated into future structure activity relationship analyses to enrich or lessen this new feature of honokiol. Two additional predicted dipyridamole-like compounds tested in this study include selumetinib and mitoxantrone, which did and did not sensitize breast cancer cells to statin-induced apoptosis. Selumetinib functions through an SREBP2-independent mechanism, suggesting that not only is the identification of feedback-dependent mechanisms beneficial for cancer treatment but also shows that additional feedback-independent classes of statin-sensitizers can be identified. This is particularly important as some multiple myeloma and prostate cancer cell lines have been shown to lack the feedback response.

The data presented here has important clinical implications for statins as anti-cancer agents. Despite some positive results from window-of-opportunity clinical trials in breast cancer using statins, a modest effect was seen from statins alone^18,19^. Therefore, discovery of novel therapeutic combinations will be necessary to achieve significant clinical impact. Since nelfinavir is poised for repurposing and statins have demonstrated anti-cancer activity in early-phase clinical trials^18,19,42–46^, clinical studies to further evaluate the therapeutic benefit of this combination could proceed swiftly. Furthermore, consideration of available gene and protein expression across our large collection of breast cancer cell lines identified a mesenchymal-enriched gene expression profile as highly predictive of sensitivity to all three statin+compound (dipyridamole, nelfinavir or honokiol) combinations. We further showed that *CDH1* expression levels served as a biomarker of synergistic response. This reinforces the dipyridamole-like behaviour of nelfinavir and honokiol, identified by our pharmacogenomics pipeline, and creates opportunities for biomarker-guided clinical studies. *CDH1* expression as a biomarker of predicted response to the combination of fluvastatin+nelfinavir could be used to identify those patients most likely to benefit. We also observed this synergistic response to the combination therapies across multiple subtypes of breast cancer. Previously we had identified the basal-like breast cancer subtype as more sensitive to statins alone; here, we have expanded the scope of statin treatment to encompass the wider breast cancer population. These findings can also be explored beyond breast cancer as *CDH1* is expressed in most cancers, for example sarcomas which are fixed in a mesenchymal state and have previously been reported as responsive to statins as single agents^37,47^.

Taken together, our computational pharmacogenomics pipeline reveals that starting with compounds that act within or on a specific pathway, it is possible to identify additional compounds to increase a class of inhibitors and/or better help understand compound mechanism of action. Our study also provides a strong preclinical rationale to warrant further investigation of the fluvastatin+nelfinavir combination, as well as the *CDH1* biomarker (**Figure 5E**). The ready availability of these well-tolerated drugs as well as simple methods for assessing *CDH1* expression could enable rapid translation of these findings to improve breast cancer outcomes.

## Methods

Our analysis design encompasses both computational identification and refinement of dipyridamole-like compounds, as well as experimental validation of the most promising candidates.

### MVA-specific Drug Network Fusion (MVA-DNF)

We developed a computational pharmacogenomic pipeline (MVA-DNF) that facilitates identification of analogues to dipyridamole, by elucidating drug-drug relationships specific to the mevalonate (MVA) pathway. MVA-DNF briefly extends upon some principles of the drug network fusion algorithm we had described previously^48^, by utilizing the similarity network fusion algorithm across three drug taxonomies (drug structures, drug perturbation, and drug sensitivity). Drug structure annotations and drug perturbation signatures are obtained from the LINCS-L1000 dataset^25,49^, and drug sensitivity signatures are obtained from the NCI-60 drug panel^26^. Drug structure annotations were converted into drug similarity matrices by calculating tanimoto similarity measures^50^ and extended connectivity fingerprints^51^ across all compounds, as described previously^48^. We extracted calculated Z-scores from drug-dose response curves for the NCI-60 drug sensitivity profiles, and computed Pearson correlation across these profiles to generate a drug similarity matrix based on sensitivity^26^. We used our PharmacoGx package (version 1.6.1) to compute drug perturbation signatures for the L1000 dataset using a linear regression model, as described previously^52^. The regression model adjusts for cell specific differences, batch effects and experiment duration, to generate a signature for the effect of drug concentration on the transcriptional state of a cell. This facilitates identification of gene expression which has been significantly perturbed due to drug treatment. These signatures indicate transcriptional changes that are induced by compounds on cancer cell lines. We further refined the drug perturbation profiles to a set of six MVA-pathway genes (**Supplementary Figure 1A**) that had been obtained from the literature as well as repositories of pathway-specific gene sets including MSigDB^53^, HumanCyc^54^ and KEGG^49,55^. These gene sets include ‘mevalonate pathway’ and ‘superpathway of geranylgeranyldiphosphate biosynthesis I (via mevalonate)’ from the HumanCyc^56^, and ‘Kegg Terpenoid Backbone Biosynthesis’ from KEGG^55,57^. The filtered drug-induced gene perturbation signatures were subsequently used to generate a drug perturbation similarity matrix that elucidates drug-drug relationships based on common transcriptional changes across the six MVA-pathway genes. We calculated similarity between estimated standardized coefficients of drug perturbation signatures using the Pearson correlation coefficient. Finally, we used the similarity network fusion algorithm^58^ to integrate drug structure, drug sensitivity, and MVA-pathway specific drug perturbation profiles, to generate an MVA-pathway specific drug taxonomy (MVA-DNF) spanning 238 compounds.

### Identification of analogues to dipyridamole

We interrogated the MVA-DNF taxonomy using a variety of approaches to identify a candidate set of dipyridamole-like compounds. Using MVA-DNF similarity scores, we first generated a ranking of all compounds closest to dipyridamole. We then conducted a perturbation test, to assess the statistical relationship of each ranked drug against dipyridamole. Briefly, drug fusion networks were generated 1000 times across perturbation, sensitivity, and drug structure profiles, each time using a random set of six genes to generate a ‘pathway-centric’ drug perturbation similarity matrix. Z-scores and p-values were calculated to determine the statistical relevance of a given dipyridamole-like analog in MVA-DNF, compared to the randomly generated networks. From this, we further ranked a list of dipyridamole-like candidate compounds by their statistical significance within MVA-DNF (p-value<0.05), resulting in identification of 23 candidate dipyridamole analogs.

For each of the dipyridamole analogues we identified, we conducted a similar assessment of significance to identify the relationships of these compounds to dipyridamole and to themselves. A network of dipyridamole-like analogues was rendered using iGraph R package^59^. Using MVA-DNF similarity scores, we further computed the contribution of each of the drug layers (structure, sensitivity and perturbation) in the identification of dipyridamole-like compounds.

We assessed the regulation of gene expression for genes involved in the mevalonate pathway across all of the top-selected dipyridamole analogues, by analyzing the drug-induced transcriptional profiles (described above) of the selected analogues. To prioritize the dipyridamole analogues, the candidate compounds were categorized, and compounds that were known toxins or carcinogens were excluded from the analysis (**Supplemental Table 1, Supplemental Figure 1C**). Top hits from the largest categories were selected for further validation.

### Cell culture and compounds

All cell lines were cultured as described previously^16,24^. Briefly, MDA-MB-231 and HCC1937 cells were cultured in Dulbecco’s Modified Eagle’s Medium (DMEM) and Roswell Park Memorial Institute medium (RPMI), respectively. All media was supplemented with 10% fetal bovine serum (FBS), 100 units/mL penicillin and 100 μg/mL streptomycin. Cell lines were routinely confirmed to be mycoplasma-free using the MycoAlert Mycoplasma Detection Kit (Lonza), and their authenticity was verified by short-tandem repeat (STR) profiling at The Centre for Applied Genomics (Toronto, ON, Canada). Fluvastatin (US Biological F5277-76) was dissolved in ethanol and dipyridamole (Sigma), nelfinavir (Sigma), honokiol (Sigma), mitoxantrone (Sigma) and selumetinib (Selleckchem) were dissolved in DMSO.

### Breast cancer cell lines panel

The breast cancer cell line^34^ panel was a generous gift from Dr. Benjamin Neel. RNAseq quantification was done using Kallisto pipeline^60^ using human transcriptome reference hg38.gencodeV23^61^. RPPA processed data was downloaded from^34^. SCMOD2^62^ breast cancer subtypes of these cell lines were obtained using genefu R package^63^.

### Breast cell-line combination viability screen

We used the sulforhodamine B colorimetric (SRB) proliferation assay^64^ in 96-well plates to determine the dose-response curves. To test the combinations in the panel of BC cell lines (See Breast cancer cell lines panel), the fluvastatin/dipyridamole, fluvastatin/nelfinavir and fluvastatin/honokiol drug combinations were tested in a 6×10 dose matrix format covering a range of decreasing concentrations of each drug (highest drug dose was 20 μM fluvastatin, 20 μM dipyridamole, 10 μM nelfinavir and 20 μM honokiol), along with all their pairwise combinations, as well as the negative control (EtOH and DMSO). We subtracted the average phosphate-buffer saline (PBS) wells value from all wells and computed the standard deviation and coefficient for each replicate. All individually treated well values were normalized to the control well values. We used Prism (v8.2.0, GraphPad Software) to compute dose-response curves with a bottom constraint equal to 0.

### Cell viability assays

3-(4,5-dimethylthiazol-2-yl)-2,5-diphenyltetrazolium bromide (MTT) assays were performed as previously described^6^. Briefly, BC cells were seeded in 750-15,000 cells/well in 96-well plates overnight, then treated in triplicate with 0-400 μM fluvastatin for 72 hours. Half-maximal inhibitory concentrations (IC_50_) values were computed from dose-response curves using Prism (v8.2.0, GraphPad Software) with a bottom constraint equal to 0.

### Cell death assays

Cells were seeded at 2.5×10^5^ cells/plates and treated the next day as indicated. After 72 hours, cells were fixed in 70% ethanol for >24 h, stained with propidium iodide and analyzed by flow cytometry for the subdiploid (% pre-G1) DNA population as a measure of cell death as previously described^6^.

### Immunoblotting

Cell lysates were prepared by washing cells twice with cold PBS and lysing cells in RIPA buffer (50 mM Tris-HCl pH 8.0, 150 mM NaCl, 0.5% sodium deoxycholate, 1% NP-40, 0.1% SDS, 1 mM EDTA, protease inhibitors) on ice for 30 min. Lysates were cleared by centrifugation and protein concentrations were determined using the Pierce 660 nm Protein Assay Kit (Thermo Fisher Scientific). Equal amounts of protein were diluted in Laemmli sample buffer, boiled for 5 min and resolved by SDS-polyacrylamide gel electrophoresis. The resolved proteins were then transferred onto nitrocellulose membranes. Membranes were then blocked for 1 hr in 5% milk in tris-buffered saline/0.1 % Tween-20 (TBS-T) at room temperature, then probed with the following primary antibodies in 5% milk/TBS-T overnight at 4 °C: SREBP-2 (1:250; BD Biosciences, 557037), p44/42 MAPK (ERK1/2) (1:1000, Cell Signaling Technology, 4695), PARP (1:1000, Cell Signaling Technology, 9542L), α-Tubulin (1:3000, Calbiochem, CP06) and E-cadherin (1:1000, Cell Signaling Technology, 3195). Primary antibodies were detected using IRDye-conjugated secondary antibodies and the Odyssey Classic Imaging System (LI-COR Biosciences). Densitometric analysis was performed using ImageJ 1.47v software.

### RNA expression analyses

Total RNA was harvested from sub-confluent cells using TRIzol Reagent (Invitrogen). cDNA was synthesized from 500 ng RNA using SuperScript III (Invitrogen). Quantitative reverse transcription PCR (qRT-PCR) was performed using the ABI Prism 7900HT sequence detection system and TaqMan probes (Applied Biosystems) for *HMGCR* (Hs00168352), *HMGCS1* (Hs00266810), *INSIG1* (Hs01650979) and *RPL13A* (Hs01578913).

### Drug combinations synergy analysis

Viability scores were calculated using standard pipelines from PharmacoGx R package^52^ and synergy scores represented by Bliss Index were calculated using SynergyFinder R package ^33^. Pearson correlation coefficient was used to measure the associations between the transcriptomic and proteomic states of cell lines and the corresponding synergy scores for each of the combinations. The transcriptomic associations were then used to rank genes for GSEA^53^. Hallmark gene set collection^35^ was downloaded from MSigDB^65^. Piano R package was used to run GSEA analysis^66^. Other EMT related pathways, namely “GO Positive Regulation of Epithelial To Mesenchymal Transition”^67^, “GO Epithelial To Mesenchymal Transition”^67^, “SARRIO Epithelial Mesenchymal Transition DN”^68^, and “SARRIO Epithelial Mesenchymal Transition Up”^68^, were also downloaded from MSigDB.

## Conflict of Interest Statement

DWC serves as a consultant for Agendia, Dynamo Therapeutics, AstraZeneca, Exact Sciences, GSK, Merck, Novartis, Pfizer, Puma, Roche; receives research support (to institution) from GSK, Merck and Pfizer and Roche-Genentech, and holds intellectual property as co-inventor on a patent related to biomarkers for TTK inhibitors. All authors declare that they have no conflicts of interest.

## Acknowledgements

This study was conducted with the support of the Terry Fox Research Institute-New Frontiers Program Project Grant (1064; LZP, BHK, WB and DWC), the Canada Research Chairs Program (to LZP; 950-229872) and Canadian Institutes of Health Research (to LZP; MOP-142263). This work was also supported by the Office of the Assistant Secretary of Defense of Health Affairs, through the Breast Cancer Research Program under Award No. W81XWH-16-1-0068 (to LZP and DWC). Opinions, interpretations, conclusions and recommendations are those of the author and are not necessarily endorsed by the Department of Defense. The authors thank all members of the Penn lab for helpful suggestions and critical feedback.

---

**Fig. S1, related to Fig. 1.**
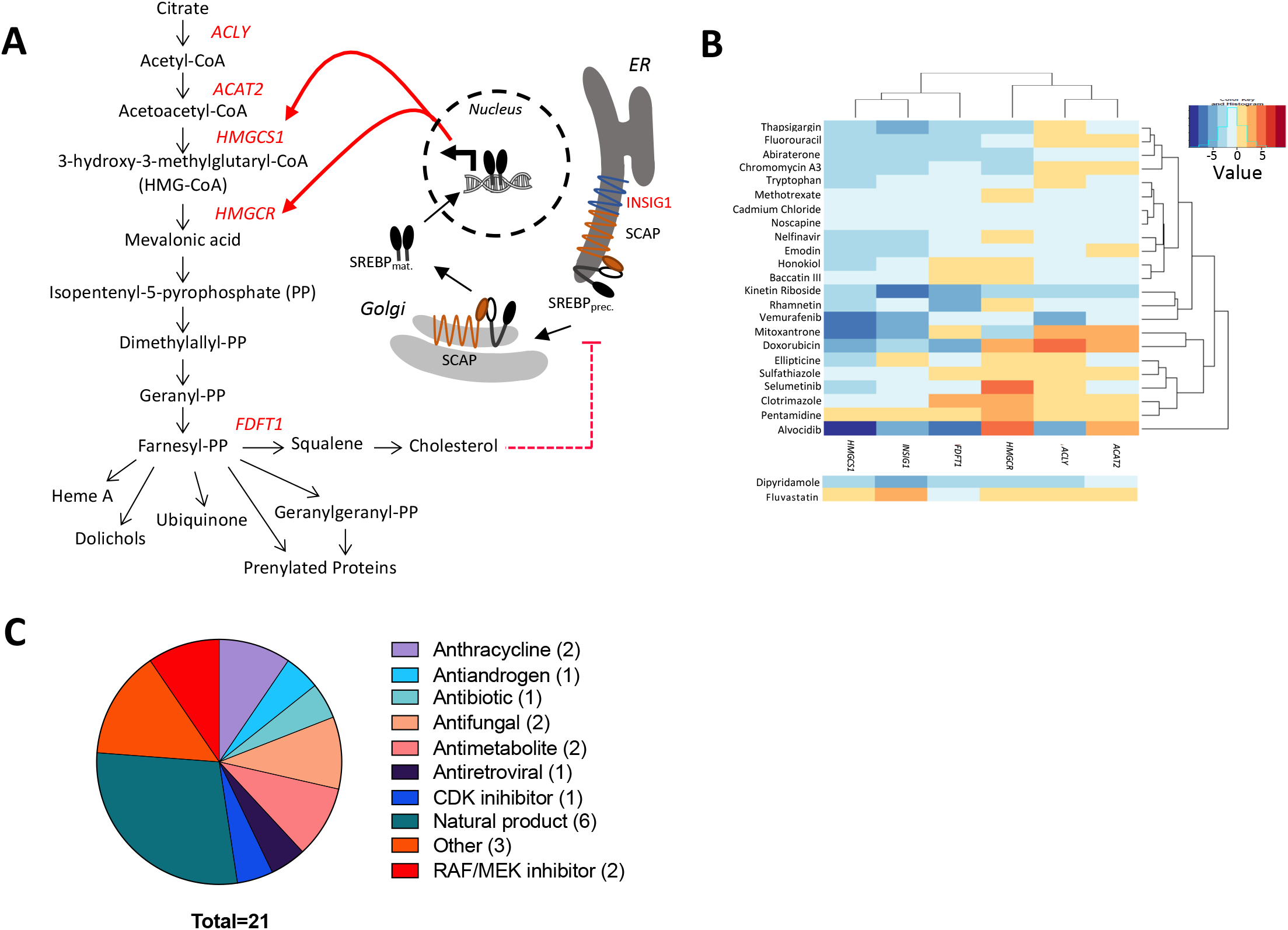
Additional information regarding drug-induced genotype changes and categorization of top 23 dipyridamole-like compounds. **(A)** Simplified schematic of the MVA pathway, highlighting the six MVA-pathway genes (in red) in the L1000 database used to restrict the drug-induced gene perturbation layer of the DNF method. **(B)** Drug perturbation signatures for dipyridamole and dipyridamole-like compounds, plotted for genes pertaining to the MVA pathway. Similarity between compounds based on their overall expression profiles is rendered in the dendrogram. Dipyridamole- and fluvastatin-induced changes shown on the bottom as reference. **(C)** Categorization of the top 21 dipyridamole-like compounds excluding toxins and carcinogenic compounds.

**Fig. S2.**
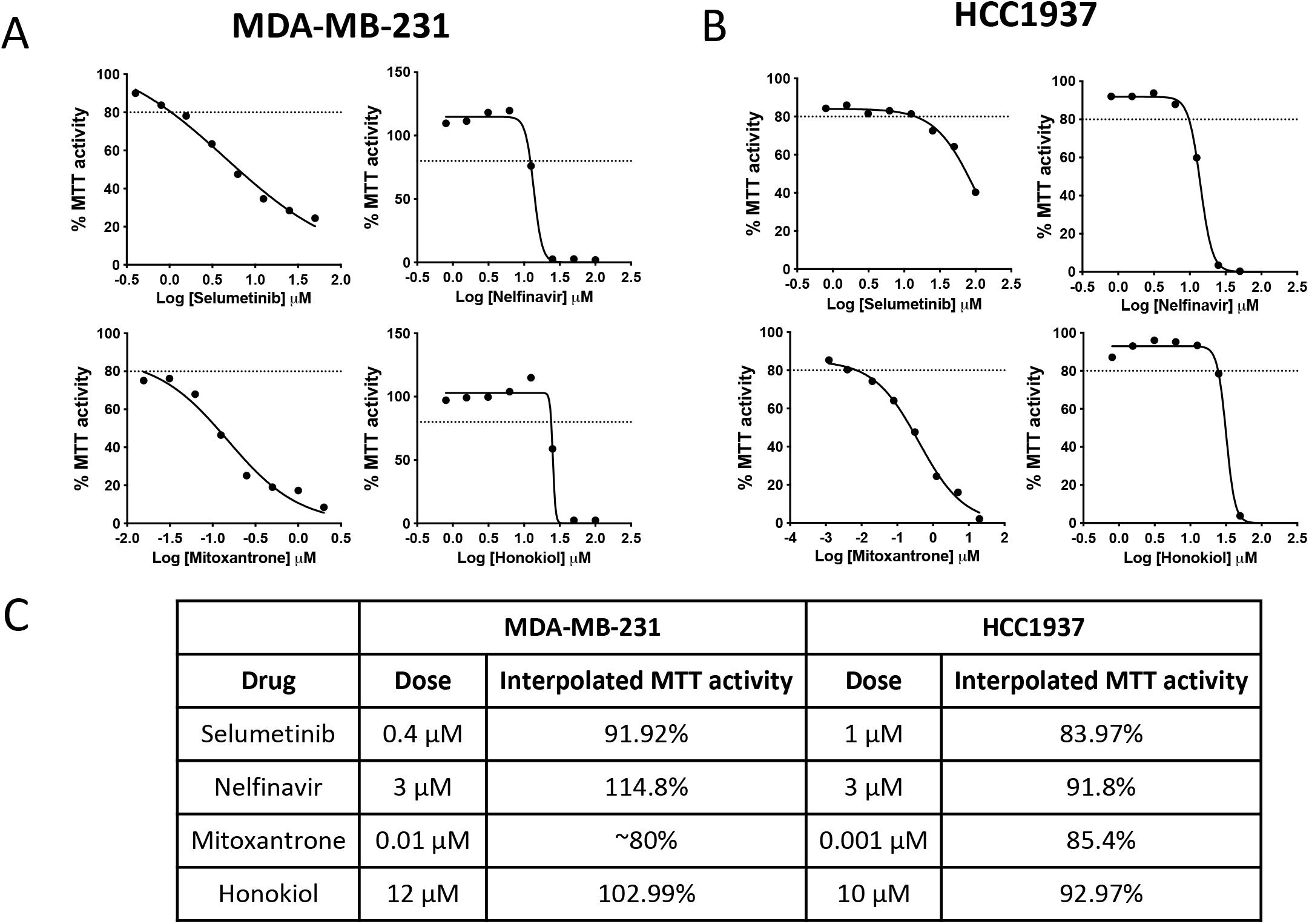
MVA-DNF drug-dose response curves for MDA-MB-231 and HCC1937 breast cancer cell lines to identify a sub-lethal dose of top dipyridamole-like compounds. **(A)** MDA-MB-231 and **(B)** HCC1937 cells were treated with a range of doses for 72 hours, and cell viability was determined using an MTT assay. The drug dose-response curves are plotted with a dashed line at 80% MTT activity indicating a sub-lethal drug dose. Data for an average of three technical replicates are plotted; data reflect the results of a single biological experiment. **(C)** Table of sub-lethal drug dose and interpolated % MTT activity for both MDA-MB-231 and HCC1937.

**Fig. S3.**
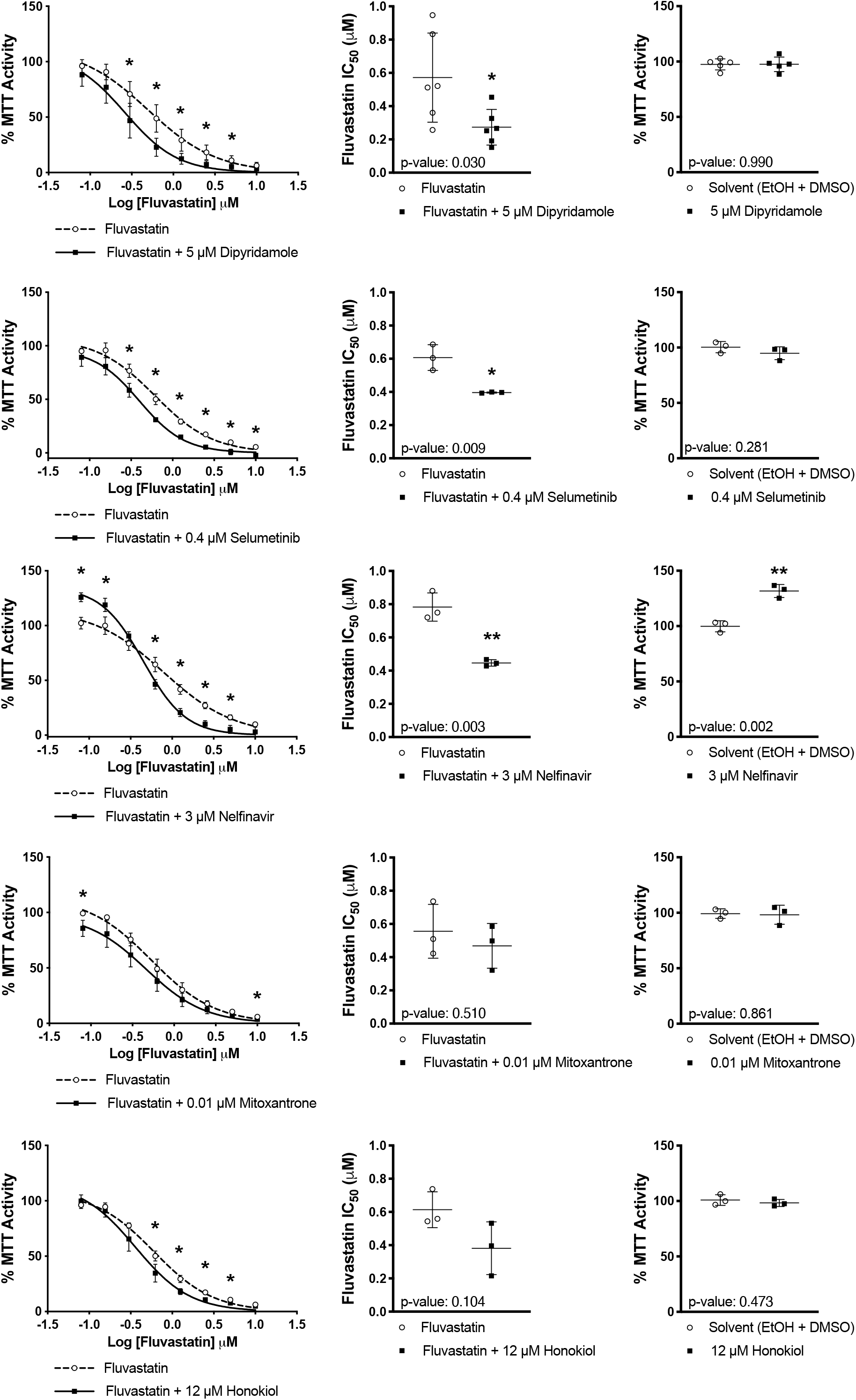
MVA-DNF drug-dose response curves, fluvastatin IC_50_ and solvent control values for MDA-MB-231 cells. MDA-MB-231 cells were treated with a range of fluvastatin doses alone or in combination with a sub-lethal dose of dipyridamole (5 μM), selumetinib (0.4 μM), nelfinavir (3 μM), mitoxantrone (0.01 μM) or honokiol (12 μM) for 72 hours, and cell viability was determined using an MTT assay. The drug dose-response curves, fluvastatin IC_50_ values and control values are plotted. Error bars represent the mean +/− SD, n = 3-5, *p <0.05, **p <0.01 (Student *t* test, unpaired, two-tailed).

**Fig. S4.**
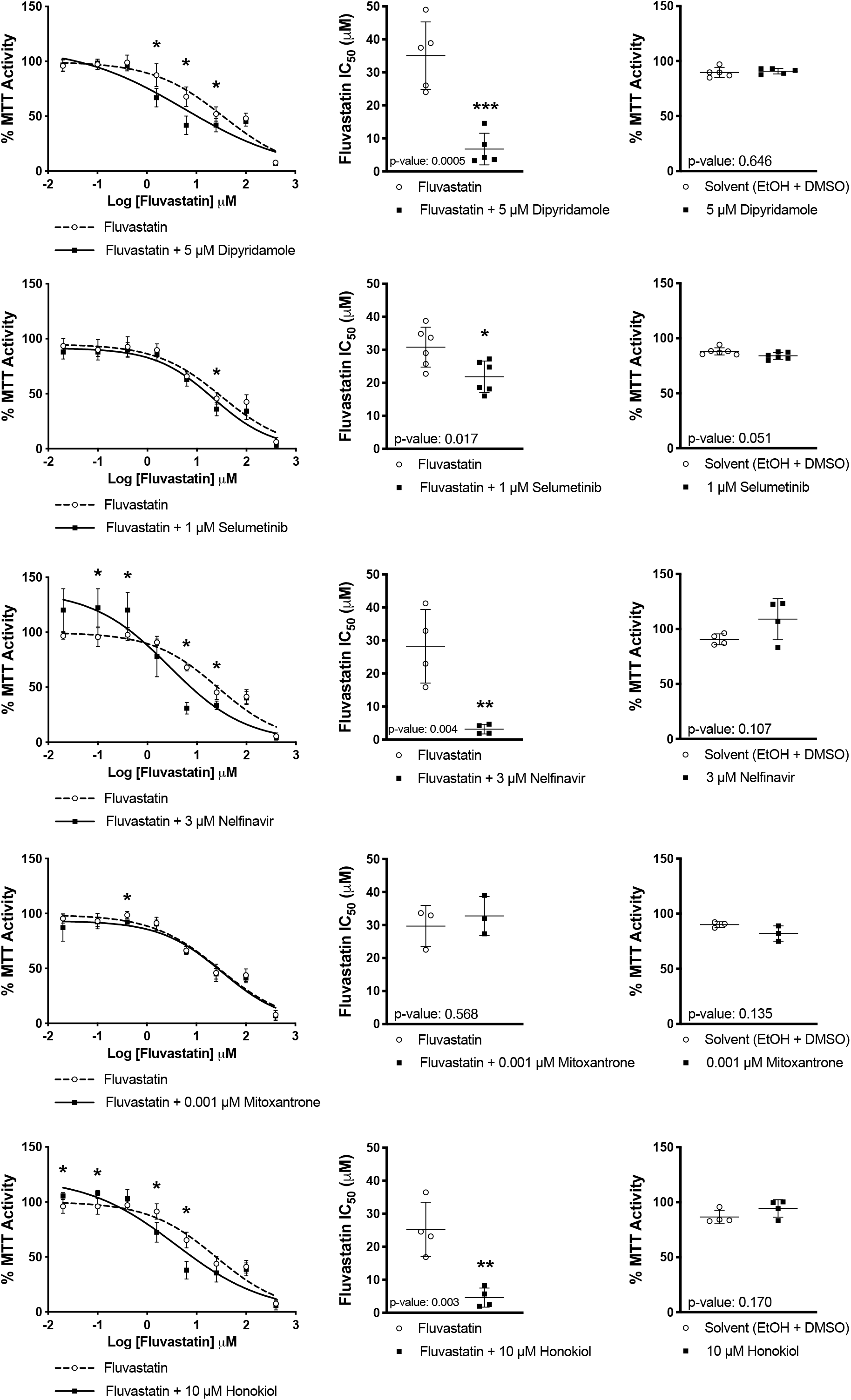
MVA-DNF drug-dose response curves, fluvastatin IC_50_ and solvent control values for HCC1937 cells. HCC1937 cells were treated with a range of fluvastatin doses alone or in combination with a sub-lethal dose of dipyridamole (5 μM), selumetinib (1 μM), nelfinavir (3 μM), mitoxantrone (0.001 μM) or honokiol (10 μM) for 72 hours, and cell viability was determined using an MTT assay. The drug doseresponse curves, fluvastatin IC_50_ values and control values are plotted. Error bars represent the mean +/− SD, n = 3-6, *p <0.05, **p <0.01, ***p <0.001 (Student *t* test, unpaired, two-tailed).

**Fig. S5, related to Fig 3.**
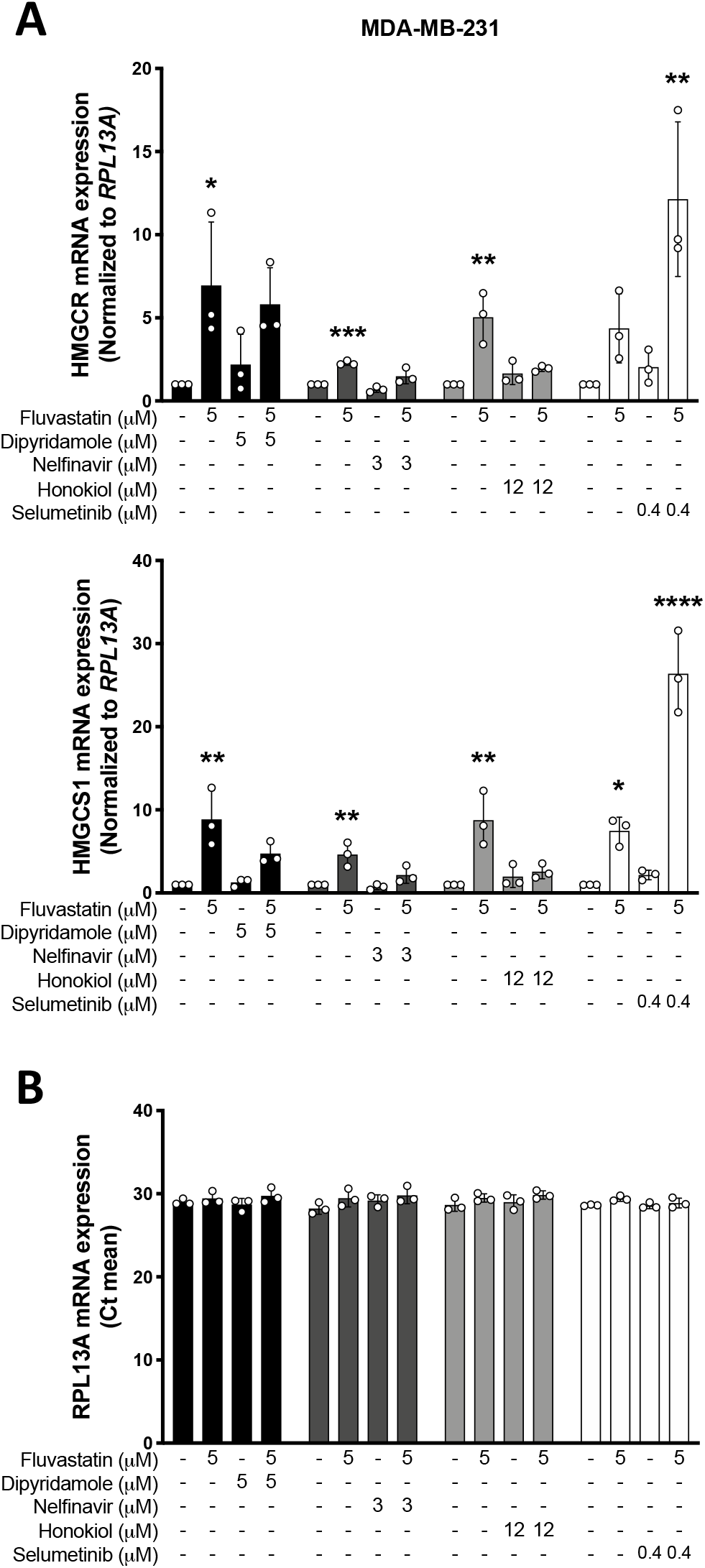
Nelfinavir and Honokiol block fluvastatin-induced SREBP activation of SREBP2 feedback genes. **(A)** MDA-MB-231 cells were treated with fluvastatin +/− dipyridamole, nelfinavir, honokiol or selumetinib for 16 hours, and RNA was isolated to assay for *HMGCR* and *HMGCS1* expression by qRT-PCR. mRNA expression data are normalized to *RPL13A* expression. **(B)** RPL13A Ct mean values plotted as a control. Error bars represent the mean +/− SD, n = 3-4, *p < 0.05, **p<0.005, ***p<0.001, ****p<0.0001 (one-way ANOVA with Bonferroni’s multiple comparisons test, where each group was compared to the solvent controls group).

**Fig. S6, related to Fig 4.**
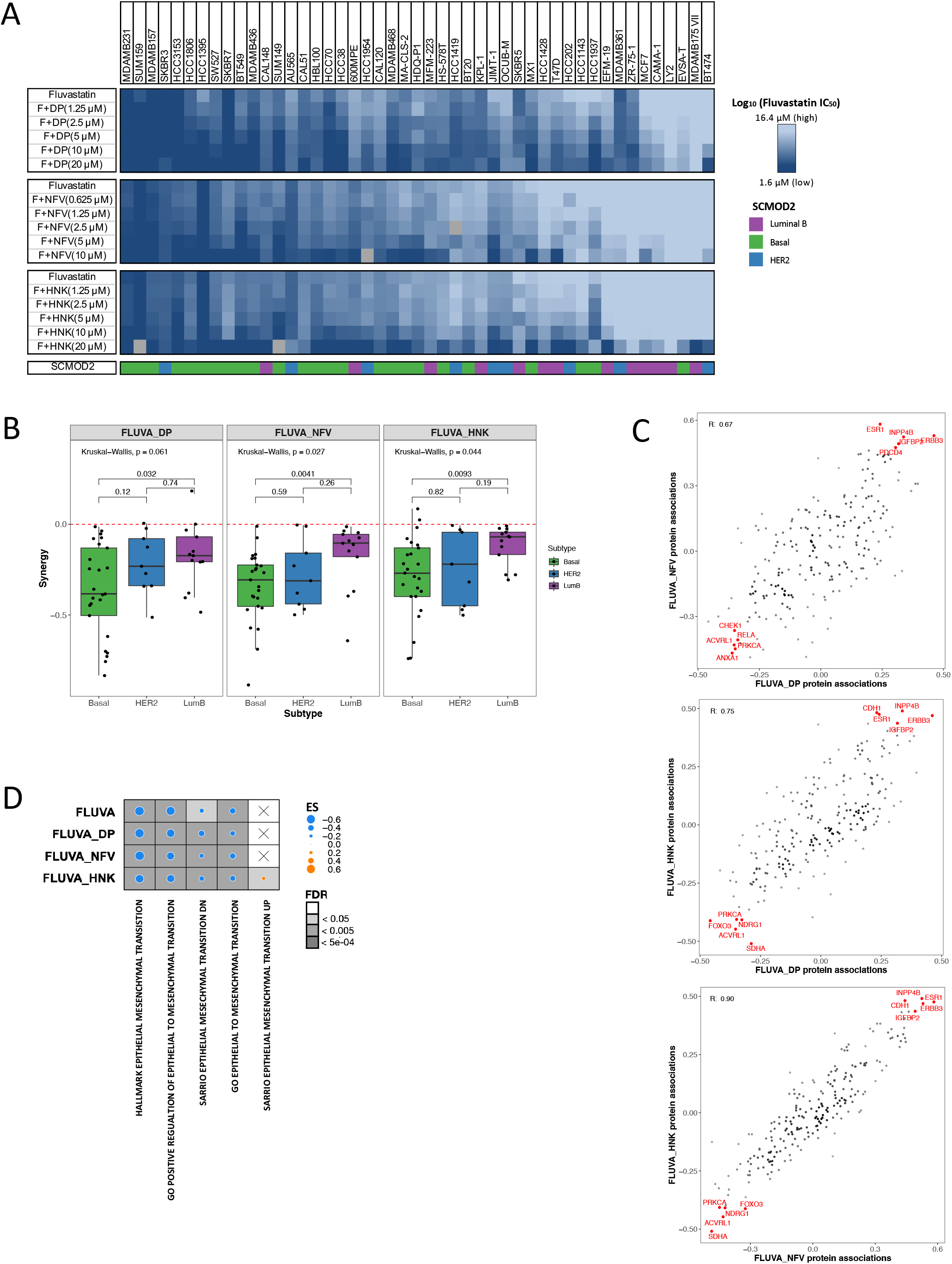
High-throughput compound combination screen. **(A)** Heatmap of Log_10_(Fluvastatin IC_50_) values for a high-throughput compound synergy screen against 47 BC cell lines visualizing the 15^th^ to 85^th^ percentile. BC cell lines were treated with a dose matrix of fluvastatin (0-20 μM) +/− dipyridamole (DP) (0-20 μM), nelfinavir (NFV) (0-10 μM) or honokiol (HNK) (0-20 μM). After 5 days of drug treatment, cell viability was assessed by SRB assay. SCMOD2 cell line subtyping was assigned to the BC cell line panel. Data presented are the average of 2 biological replicates (fluvastatin +/− dipyridamole (DP)) or the mean of 3-6 biological replicates (fluvastatin +/− nelfinavir (NFV) and fluvastatin +/− honokiol (HNK)). **(B)** Comparison of synergy scores stratified by BC subtypes across the combinations using wilcoxon paired rank test. Red dash line at synergy threshold. **(C)** Similarity of proteomic states associations^34^ with synergy scores across the fluvastatin+compound combinations. Similarity of proteomic states associations were compared across the combinations (Fluva+DP vs Fluva+NFV; F+DP vs Fluva+HNK; Fluva+NFV vs Fluva+HNK) using Pearson correlation coefficient. Top five basally-expressed proteins associated with synergy in either direction are annotated in red. **(D)** Gene set enrichment analysis using five EMT gene set collections and genes ranked by basal mRNA correlated to the fluvastatin IC_50_ (Fluva) value or synergy score (Fluva+DP, Fluva+NFV and F+HNK). Dot size indicates the difference in enrichment scores (ES) of the pathways. Background shading indicates the FDR. X indicates pathway and drug combinations that were not significantly enriched (FDR > 0.05).

**Fig. S7, related to Fig 4.**
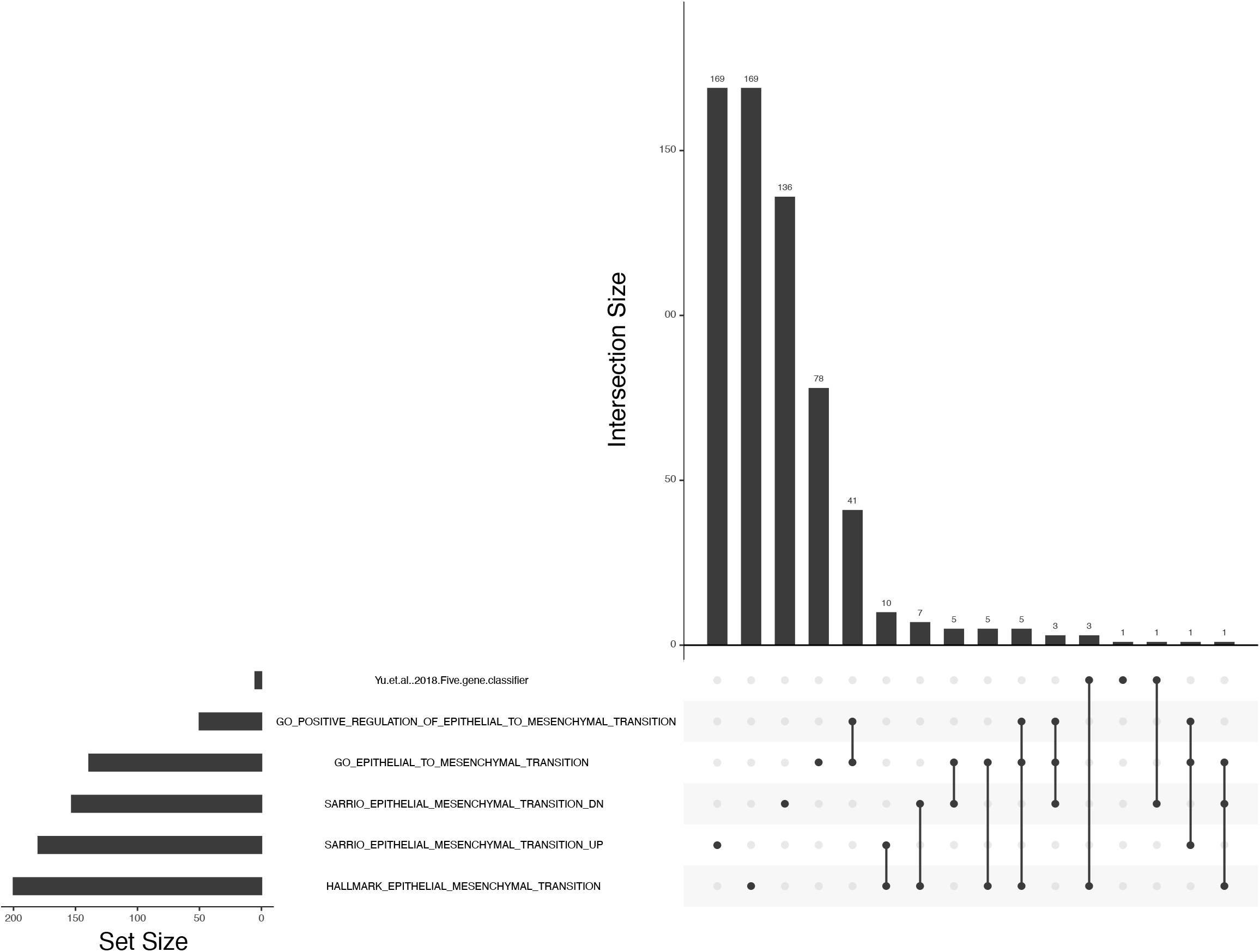
Overlapping genes within the EMT gene sets. **(A)** Upset plot to visualize the agreement between Yu *et al.* (2017)^36^ five-gene classifier and five additional EMT gene sets.

**Fig. S8, related to Fig 5.**
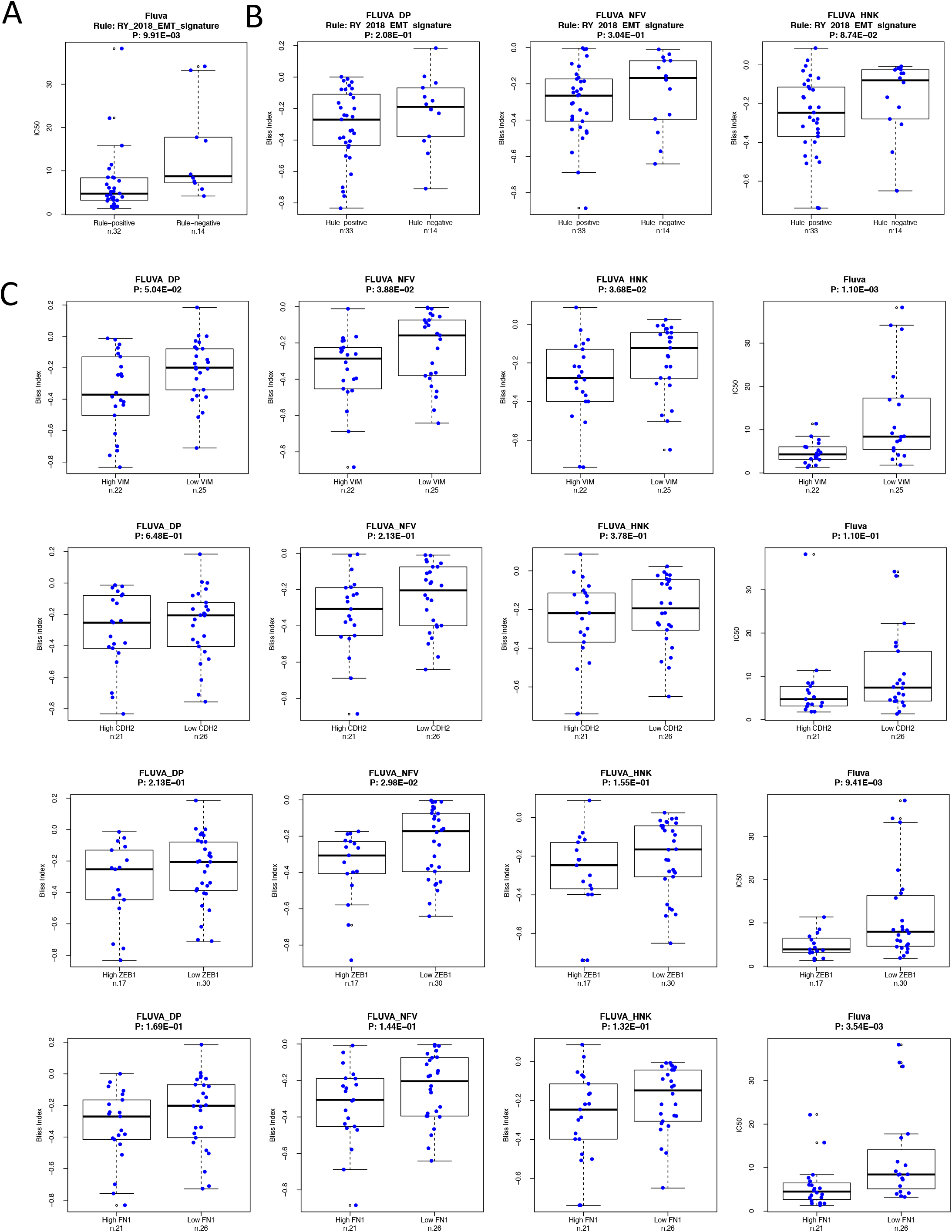
EMT gene expression as a biomarker of sensitivity to fluvastatin and synergistic response to fluvastatin+compound combinations. **(A)** Five-gene fluvastatin sensitivity gene classifier^36^ predicts sensitivity to fluvastatin alone, but **(B)** does not predict synergy to F+DP, F+NFV or F+HNK. **(C)** Basal Vimentin (VIM), N-Cadherin (CDH2), ZEB1 and fibronectin (FN1) mRNA expression do not predict synergy to the drug combinations.

## Table Legends

**Supplementary Table 1 -.** Ranked MVA-DNF compounds by Z-score.

| Hits | Permutation Test |  | Currently used<br>in humans | Currently used<br>in cancer<br>treatment | In clinical trials<br>for cancer<br>treatment | Pre-clinical | Research tool | Exclude | Classification |
| --- | --- | --- | --- | --- | --- | --- | --- | --- | --- |
|  | Z-Score | P-Value* |  |  |  |  |  |  |  |
| SELUMETINIB | -3.57 | 1.77E-04 | + |  | + |  |  |  | RAF/MEK inhibitor |
| NELFINAVIR | -3.28 | 5.18E-04 | + |  | + |  |  |  | Antiretroviral |
| MITOXANTRONE | -3.20 | 6.82E-04 | + | + | + |  |  |  | Anthracycline |
| DOXORUBICIN | -3.03 | 1.22E-03 | + | + | + |  |  |  | Anthracycline |
| HONOKIOL | -2.96 | 1.53E-03 |  |  |  | + |  |  | Natural product |
| CLOTRIMAZOLE | -2.95 | 1.57E-03 | + |  |  |  |  |  | Antifungal |
| SULFATHIAZOLE | -2.89 | 1.90E-03 |  |  |  | + |  |  | Antibiotic |
| VEMURAFENIB | -2.67 | 3.79E-03 | + | + | + |  |  |  | RAF/MEK inhibitor |
| CHROMOMYCINA3 | -2.64 | 4.13E-03 |  |  |  | + | + | Toxin | Antibiotic |
| BACCATINIII | -2.53 | 5.69E-03 |  |  |  | + |  |  | Natural product |
| NOSCAPINE | -2.51 | 6.11E-03 |  |  | + | + |  |  | Natural product |
| METHOTREXATE | -2.29 | 1.11E-02 | + | + | + |  |  |  | Antimetabolite |
| CADMIUMCHLORIDE | -2.22 | 1.33E-02 |  |  |  | + | + | Carcinogen | Other |
| RHAMNETIN | -2.18 | 1.45E-02 |  |  |  | + |  |  | Natural product |
| PENTAMIDINE | -2.18 | 1.46E-02 | + |  | + |  |  |  | Antifungal |
| EMODIN | -2.16 | 1.55E-02 |  |  |  | + |  |  | Natural product |
| FLUOROURACIL | -2.11 | 1.75E-02 | + | + | + |  |  |  | Antimetabolite |
| TRYPTOPHAN | -1.92 | 2.76E-02 | + |  | + |  |  |  | Other |
| ALVOCIDIB | -1.82 | 3.44E-02 | + | + | + |  |  |  | CDK inhibitor |
| ABIRATERONE | -1.77 | 3.81E-02 | + | + | + |  |  |  | Antiandrogen |
| THAPSIGARGIN | -1.71 | 4.36E-02 |  |  |  | + |  |  | Other |
| KINETINRIBOSIDE | -1.66 | 4.84E-02 |  |  |  | + |  |  | Other |
| ELLIPTICINE | -1.65 | 4.92E-02 |  |  |  | + |  |  | Natural product |

\* Table ordered by p-value

